# Genomic and microscopic evidence of stable high density and maternally inherited *Wolbachia* infections in *Anopheles* mosquitoes

**DOI:** 10.1101/2020.10.29.357400

**Authors:** Thomas Walker, Shannon Quek, Claire L. Jeffries, Janvier Bandibabone, Vishaal Dhokiya, Roland Bamou, Mojca Kristan, Louisa A. Messenger, Alexandra Gidley, Emily A. Hornett, Enyia R. Anderson, Cintia Cansado-Utrilla, Shivanand Hegde, Chimanuka Bantuzeko, Jennifer C. Stevenson, Neil F. Lobo, Simon C. Wagstaff, Christophe Antonio Nkondjio, Eva Heinz, Grant L. Hughes

## Abstract

*Wolbachia*, a widespread bacterium that can reduce pathogen transmission in mosquitoes, has been detected within populations of *Anopheles (An.)* malaria vectors. In the *An. gambiae* complex, the primary vectors in Sub-Saharan Africa, *Wolbachia* strains are at low density and infection frequencies in wild populations. PCR-independent evidence is required to determine whether *Wolbachia* strains are true endosymbionts in *Anopheles* given most studies to date have used nested-PCR to identify strains. Here we report high-density strains found in geographically diverse populations of *An. moucheti* and *An*. *demeilloni*. Fluorescent *in situ* hybridization localized a heavy infection in the ovaries of *An. moucheti* and maternal transmission was observed. Genome sequencing of both strains obtained genome depths and coverages comparable to other known infections. Notably, homologs of cytoplasmic incompatibility factor (*cif*) genes were present indicating these strains possess the capacity to induce the phenotype cytoplasmic incompatibility which allows *Wolbachia* to spread through populations. The characteristics of these two strains suggest they are ideal candidates for *Wolbachia* biocontrol strategies in *Anopheles*.

## Introduction

The endosymbiotic bacterium *Wolbachia* is currently being deployed into the field for population replacement and population suppression vector control strategies. These approaches are showing great promise in *Aedes (Ae.)* mosquitoes, particularly *Ae. aegypti*^1–5^ which is the main vector of arboviruses such as dengue, Chikungunya and Zika viruses. However, translating the control strategy into *Anopheles* mosquitoes for malaria control is proving more challenging, mainly due to the inability to create stable *Wolbachia* transinfected lines in the major mosquito vectors from sub-Saharan Africa. The development of novel malaria control tools is highly desirable as the emergence of insecticide resistance impacts the effectiveness of current control strategies^6^.

*Wolbachia* can induce two desirable properties in mosquitoes that are exploited for vector control; inhibition of pathogens and cytoplasmic incompatibility (CI). Although the application of *Wolbachia*-based biocontrol for malaria is still in its infancy, there is evidence that it could emulate the success seen for arboviruses if stable lines are developed. For example, experiments investigating both transient *Wolbachia* infections in *An. gambiae*^7^ and stable transinfected *Wolbachia* in *An. stephensi*^8^ demonstrated a reduction in the density of *Plasmodium (P.) falciparum* malaria parasites. The level of inhibition of *Plasmodium* parasites is dependent on the particular strain of *Wolbachia*^9–11^. In the stable transinfected *An. stephensi* line, *Wolbachia* was able to spread into caged populations by CI, although repeated male introductions were required, indicating there was a fitness cost associated with the infection^8,12^. These studies show that pathogen inhibition and CI is induced in *Anopheles* although optimal strains are likely required before translation to the field can become a reality. Strains derived from native *Wolbachia* infections in *Anopheles* species may be more effective for transinfection into other medically important vector species such as *An. gambiae* and *An. coluzzii*, as they are more likely to form mutualistic partnerships without deleterious effects on host biology. As such, it is essential to have a thorough understanding of the infection status of native *Wolbachia* strains in *Anopheles* and determine their phenotypic effects.

While the standing dogma in the *Wolbachia* field for many years was that *Anopheles* mosquitoes were impervious to *Wolbachia* infection^13,14^, there have been numerous recent reports of natural infections in a range of *Anopheles* species^15–22^. However, the majority of these studies found low density, low prevalence *Wolbachia* infections in diverse *Anopheles* species. The reliance on only a few genes (particularly *16S rRNA*) to determine the phylogeny of newly discovered strains is also problematic. By their very nature, low-density *Wolbachia* infections are challenging to confirm, and the prominent use of nested PCR to characterise these infections has led to questioning of the validity of these strains^23,24^. The detection of gene sequences does not necessarily confirm the presence of endosymbiotic bacteria given the possibility of environmental contamination or integration into the host genome^23^. Furthermore, low prevalence rates in wild mosquito populations would also indicate that these strains are unlikely to be inducing CI.

Previously we identified potentially high density *Wolbachia* infections in *An. moucheti*, *An. species* A and another unclassified *Anopheles* species suggesting these strains may be more appropriate candidates for biocontrol strategies^22,25^. Here we analysed larger cohorts of mosquitoes from Cameroon, the DRC and Kenya and *Wolbachia* prevalence rates in wild mosquito populations varied from 38-100% depending on strain and locality. Importantly, we undertook fluorescence *in situ* hybridization (FISH) which confirmed high density *Wolbachia* infection in the germline tissue. We quantified *Wolbachia* density and demonstrated that both strains infect somatic tissues and are maternally transmitted. Additionally, we performed Illumina sequencing and present the first complete genome sequences of two strains that naturally reside in *Anopheles* species. Genome coverage for both strains was >50x (up to five-fold higher prevalence than *Wolbachia* in *Ae. albopictus*) and *16S rRNA* sequencing shows that these *Wolbachia* strains exhibit dominance within the microbiome providing further evidence that these *Wolbachia* strains are high density strains that naturally infect *Anopheles* species. Taken together our results provide robust evidence for natural *Wolbachia* strains in *Anopheles* mosquitoes.

## Results

### *Wolbachia* prevalence rates

*Wolbachia* strains that induce CI and can invade mosquito populations are more likely to result in high prevalence rates in wild populations. Here we undertook a robust screening approach examining 1582 mosquitoes from Cameroon, the DRC and Kenya to determine the prevalence in wild populations and quantify vertical transmission. *Wolbachia* qPCR analysis of a large number of wild adult female *An. moucheti* from Cameroon (n=1086) revealed an overall prevalence of 56.6% for the *w*AnM strain **(Fig. 1a)** that we previously discovered in the DRC^22^. *Wolbachia* was detected in 85.7% (6/7) of *An. moucheti* adult females collected from the DRC in 2015. To determine if the intermediate prevalence rate in *An. moucheti* field populations in Cameroon could be influenced by host genetic diversity, we first designed mosquito ITS2 species-specific qPCR assays to rapidly confirm species (**Extended data Fig. 1 and 2**). As we observed some high ITS2 Ct values in a small number of individual mosquitoes, sequencing of the internal transcribed spacer 2 (ITS2) region and the *cytochrome oxidase subunit II* (*COII)* gene was undertaken on a subsample and revealed the presence of two sub-groups (‘*An. m. moucheti’* and ‘*An. m. cf moucheti*’) in wild populations in Cameroon **(Fig. 1b-c, Supplementary table 1).**

**Fig. 1:**
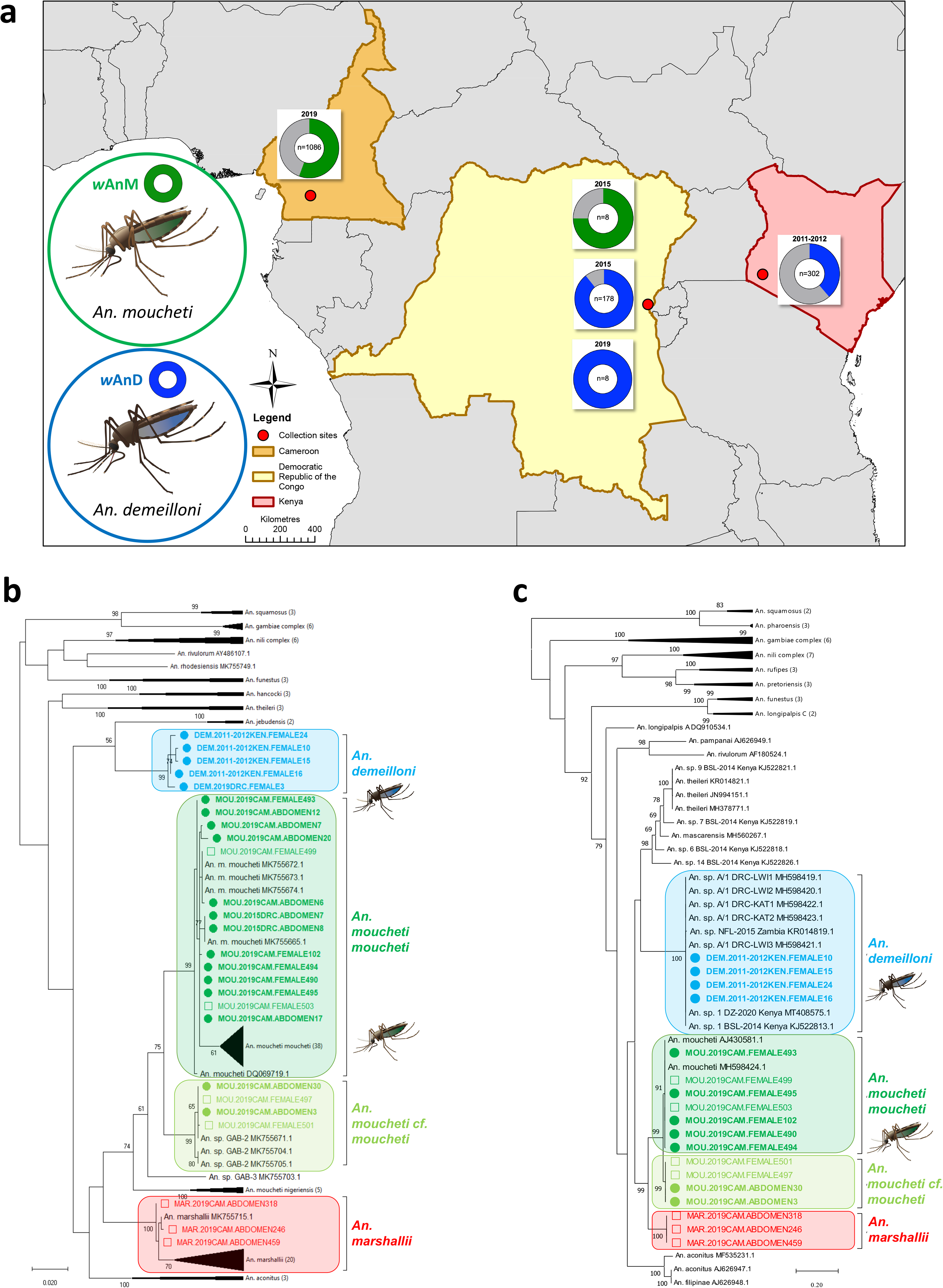
Mosquito collection sites, *Wolbachia* prevalence rates and host mosquito phylogenetic analysis. **a**, *Wolbachia* prevalence rates in wild adult female mosquitoes for the *w*AnM strain in *An. demeilloni* and *w*AnM strain in *An. moucheti* are denoted in blue and green respectively. **b,** Mosquito *COII* phylogenetic tree with the highest log likelihood (−4605.97). The analysis involved 130 nucleotide sequences with a total of 735 positions in the final dataset. Filled circles = *Wolbachia*-infected individuals, open squares = uninfected individuals. **c,** Mosquito ITS2 phylogenetic tree with the highest log likelihood (−11797.51). The analysis involved 71 nucleotide sequences. Codon positions included were 1st+2nd+3rd+Noncoding. There was a total of 1368 positions in the final dataset. Filled circles = *Wolbachia*-infected individuals, open squares = uninfected individuals. **b,c** Reference numbers of additional sequences obtained from GenBank (accession numbers) are shown unless subtree is compressed. The trees are drawn to scale, with branch lengths measured in the number of substitutions per site.

We had previously discovered a novel *Wolbachia* strain in an unidentified *Anopheles* species (referred to as *An. species* A^22^). Using morphological keys^26,27^ on freshly collected individuals we confirmed this species to be *An. demeilloni* (**Extended data Fig. 3**). We detected *Wolbachia* (now termed the *w*AnD strain) in 38.7% (117/302) of females collected from Kenya in 2011-2012. The *w*AnD strain was present in 89.3% (159/178) of females collected from the DRC in 2015 and in 100% (n=8) of females from the DRC collected in 2019. ITS2 species-specific PCR assays and sequencing with phylogenetic comparison to previously published *An. species* A sequences^22,28^ were used to confirm all individuals analysed were *An. demeilloni* (**Fig. 1b-c, Supplementary table 1**).

### *Wolbachia* strains are maternally inherited and can be visualised in mosquito ovaries

The high prevalence of *Wolbachia* compared to previous studies of other *Anopheles* species led us to speculate that vertical transmission was maintaining the bacteria in the populations at high rates. To investigate this, we attempted to obtain progeny from field-collected mosquitoes. While colonising mosquitoes is challenging, we were able to acquire progeny, although there was substantial mortality of F1 larvae in both species. We screened surviving generations and detected *w*AnM in the resulting F1 generation from Cameroon wild-caught *An. moucheti* females and *w*AnD in the F1 and F2 *An. demeilloni* generations resulting from wild-caught females from the DRC (**Fig. 2a, Supplementary table 2**). In both *An. moucheti* and *An. demeilloni* we detected *Wolbachia* in all developmental stages. We were able to maintain the *An. demeilloni* colony to the F2 generation and detected *Wolbachia* in the adult female abdomens (10/11) and in the head and thorax (2/11).

**Fig. 2:**
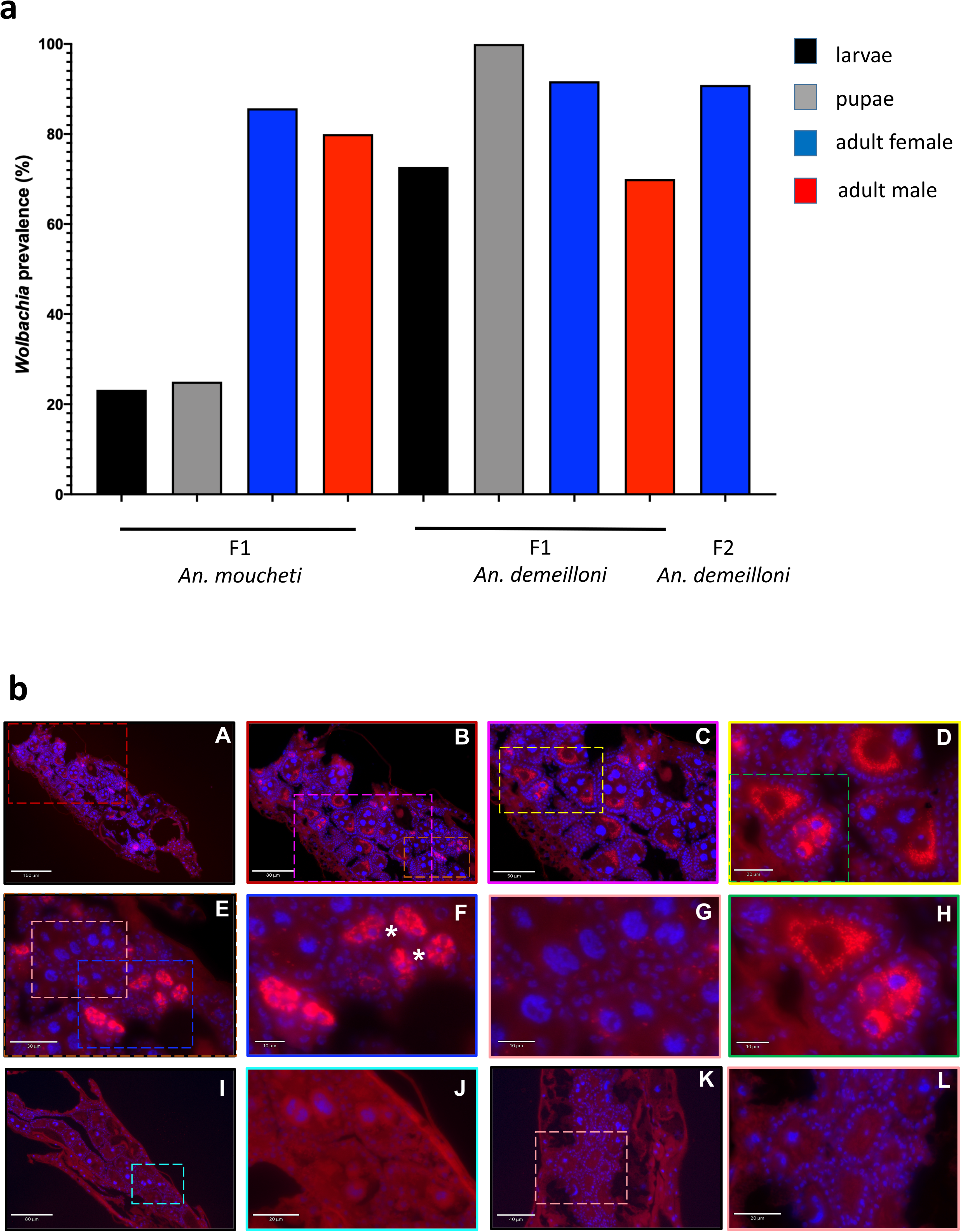
Maternal transmission and visualisation of *Wolbachia* in mosquito ovaries. **a**, Wild caught mosquitoes were offered a blood meal to support egg development and offered an oviposition site. Eggs were hatched and the prevalence of vertically transmitted *Wolbachia* was determined in the F1 larvae, pupae or adults using qPCR. **b,***Wolbachia* was primarily located to the ovarian follicles (A-H). Coloured boxes indicate area of magnification for subsequent images. Within the same ovary, some ovarian follicles are sparsely infected with *Wolbachia* (E and magnification in G) while others have a heavy infection (C, D and H; E and F). Asterisks indicate infection in the secondary follicles. *Wolbachia* was imaged with an Alexa 590 labelled probe targeting the *Wolbachia 16S rRNA* gene (red) and DNA was stained with DAPI (blue). No probe control images (I – L) show no fluorescent signal (I & J and K & L are two separate individuals). FISH analysis found nine out of 16 individuals positive for infection.

Several recent studies have called for microscopy to validate PCR data when determining the presence of *Wolbachia* strains in wild mosquito populations^23,24^. As such we undertook FISH to visualise *Wolbachia* in *An. moucheti*. *Wolbachia* could be clearly seen in the mosquito ovaries of 9/16 (56.3%) wild-caught females with a heavy infection observed in the ovarian egg chambers (**Fig. 2b, Extended data Fig. 4)**. Confocal microscopy was used to further resolve the location of the infection with *Wolbachia* observed within the oocyte surrounding the nuclei. Lower levels of *Wolbachia* were seen in the nurse cells. This infection pattern was similar to transinfected stable lines generated in *An. stephensi*^8^, although there appears a lower bacterial density in the germline in the transinfected lines compared to the native *w*AnM strain in *An. moucheti*. It was noticeable that some ovarian follicles had a heavy *w*AnM infection while for others the infection was sparse. While the reasons for the differences in loading of the ovarian follicles remains to be determined this may explain the heterogenous infection prevalence we found in field populations.

### *Wolbachia* strains are high density and infect somatic tissues

The majority of studies that have identified *Wolbachia* in *Anopheles* species have used nested PCR indicating low density infections. We previously presented evidence that *w*AnM and *w*AnD are likely significantly higher-density strains^22^ and our microscopy from this study shows a high density of *Wolbachia* bacteria in *An. moucheti* ovaries. We further characterised density using qPCR on large cohorts of wild-caught females and showed significant variation in *Wolbachia* density across mosquito species, body parts and life cycle stages (**Supplementary table 2).** When comparing the density of *w*AnM in the abdomen (n=377) and corresponding head-thorax (n=99) of *An. moucheti* wild-caught females from Cameroon, the density was significantly higher in abdomen extractions (Students t-test, p=0.0003) (**Fig. 3a**). Interestingly, we found a significantly higher *w*AnD density in *An. demeilloni* adult females collected from Kenya (n=117) compared to the DRC in 2015 (n=158) (Students t-test, p<0.0001) (**Fig. 3a**). When comparing the overall *Wolbachia* densities in whole-body females, the *w*AnM strain in *An. moucheti* from Cameroon collected in 2019 (n=238) was significantly higher compared to the combined cohorts of *w*AnD strain in *An. demeilloni* from both the DRC in 2015 and Kenya in 2011-2012 (n=402) (Students t-test, p=0.003).

**Fig. 3:**
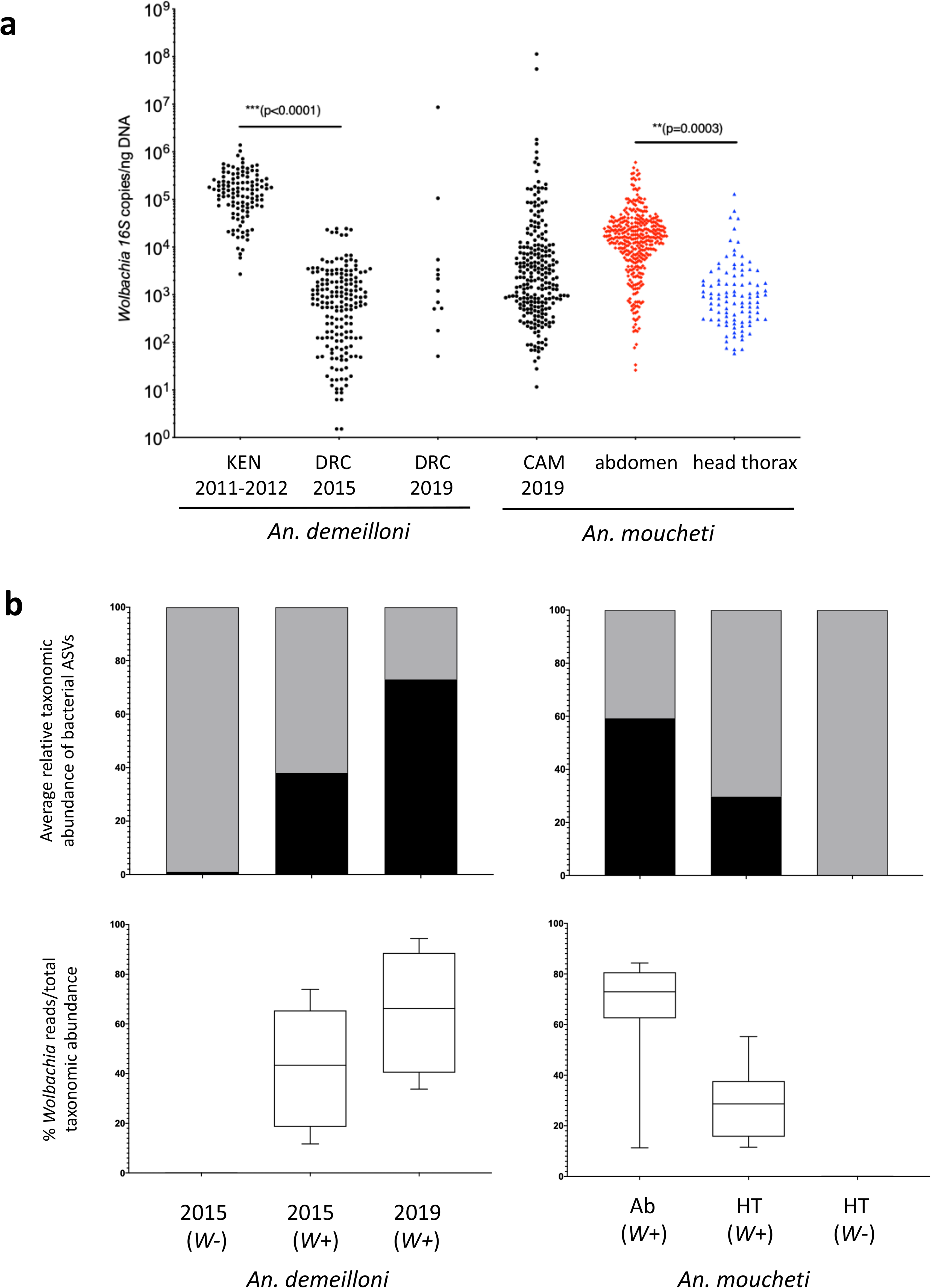
*Wolbachia* strain densities and relative abundance in the mosquito microbiome. **a,** normalised *Wolbachia* strain densities measured using qPCR of the conserved *Wolbachia 16S rRNA* gene. A synthetic oligonucleotide standard was used to calculate *Wolbachia 16S rDNA* gene copies per ng total DNA using a ten-fold serial dilution standard curve. **b,** Relative *Wolbachia* abundance in the mosquito microbiome. Average relative taxonomic abundance of bacterial ASVs within the *16S rDNA* microbiomes of *An. demeilloni* and *An. moucheti* using QIIME2^66^. *Wolbachia* is represented in black, all other bacteria are represented in grey (more information available in Extended data Fig. 5). *Wolbachia* % abundance of total *16S rDNA* bacterial load is seen through box-and-whisker plots.

### *Wolbachia* strains dominate the microbiome

To further confirm the high-density infections, we analysed the composition of bacterial species within selected *An. demeilloni* and *An. moucheti* adult females to determine the relationship of resident *w*AnD and *w*AnM strains and other bacteria (**Fig. 3b, Extended data Fig. 5**,). For wild-caught *An. demeilloni* females collected from the DRC in 2015 (n=9), *Wolbachia* is the dominant amplicon sequence variant (ASV) when present, compromising an average 38.1% of total *16S rRNA* reads. In *An. demeilloni* females collected in 2019 (n=8), *Wolbachia* reads comprise 72.6% of the microbiome. For comparison, we analysed a selection of *An. demeilloni* 2015 wild-caught females that were *Wolbachia* negative by PCR (n=6) and found no *Wolbachia* reads (**Fig. 3b**). We also examined the microbiome profiles of *An. moucheti Wolbachia*-infected abdomens, *Wolbachia*-infected head-thoraxes and *Wolbachia*-uninfected head-thoraxes from Cameroon (**Fig 3b**). Again, in our PCR-positive samples, *Wolbachia* was the dominant ASV in abdomens (average 59.2%, n=19) and in the head-thorax samples (average 29.7%, n=8). Our microbiome data corroborate our PCR results with minimal *Wolbachia* reads in our uninfected *An. moucheti* head-thorax samples (n=6).

### *Wolbachia* strain variation

Another characteristic of stably infected *Wolbachia* strains is the presence of the same strain in geographically distinct populations of the same insect species. We undertook Multilocus Sequencing Typing (MLST) and found identical allelic profiles for *w*AnM-infected *An. moucheti* (n=3) from Cameroon in comparison to the *w*AnM strain in *An. moucheti* from the DRC ^22^. Further analysis of the *Wolbachia* surface protein (*wsp)* gene (n=50) resulted in three replicates of the same variant sequence (all with the same three single nucleotide polymorphisms) (**Fig. 4**) seen within hypervariable region 2 (**Supplementary table 3**). Using mosquito *COII* gene phylogeny, the three variant *Wolbachia wsp* sequences were from *An. m. cf moucheti*, whereas the non-variants (n=47) were from *An. m. moucheti*. Analysis of *wsp* gene sequences from *w*AnD-infected *An. demeilloni* from Kenya (n=40) revealed no variation. However, MLST sequences from three individuals resulted in evidence of a *coxA* gene sequence variant previously shown in the DRC ^22^ (**Supplementary table 3**). These results provide strong evidence for the presence of the same *w*AnM and *w*AnD strain variants across large geographical areas.

**Fig. 4:**
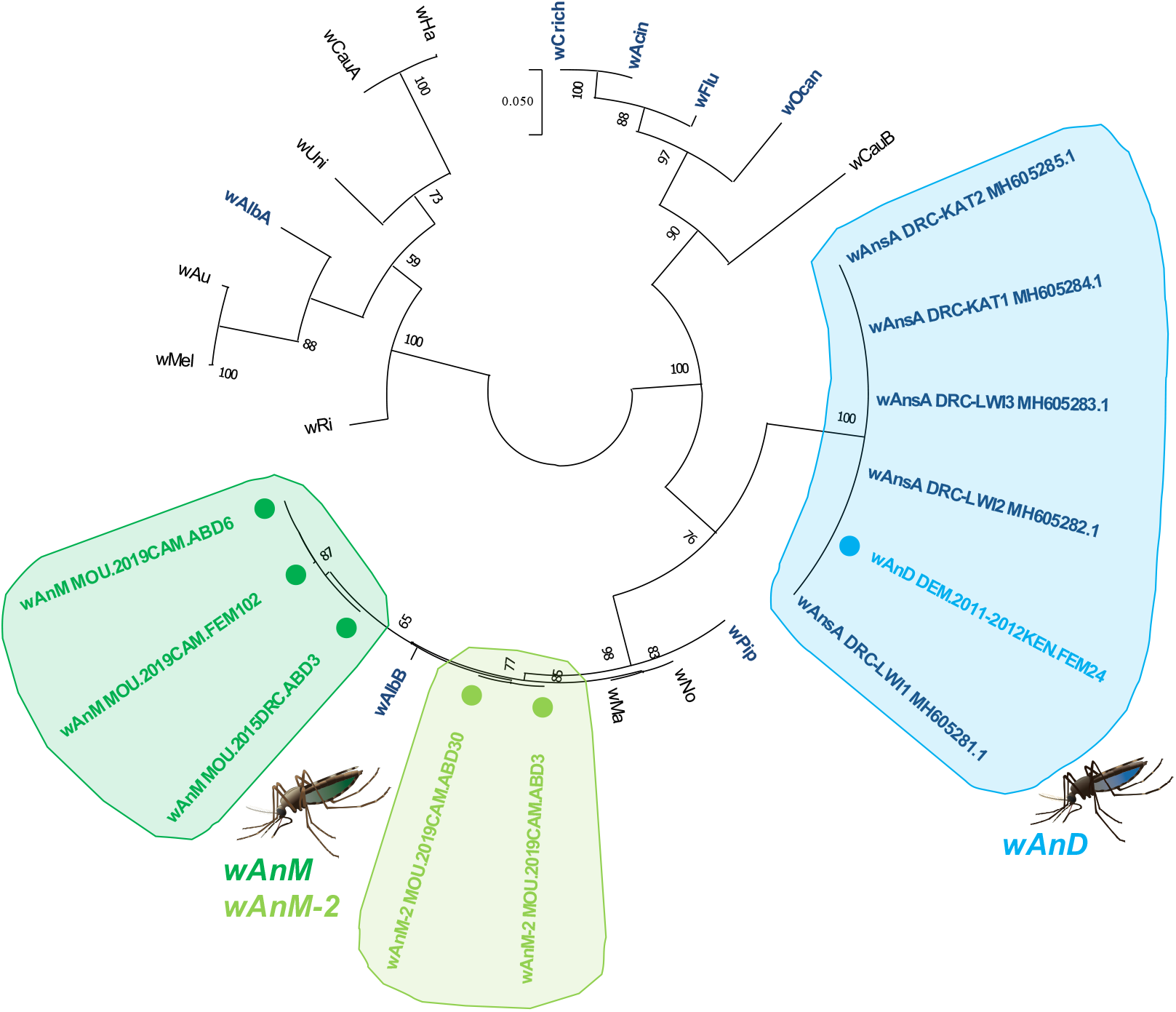
Maximum likelihood molecular phylogenetic analyses of the *Wolbachia wsp* gene. The tree with the highest log likelihood (−3004.54) is shown and the analysis involved 27 nucleotide sequences. Codon positions included were 1st+2nd+3rd+Noncoding. There was a total of 586 positions in the final dataset.

### *Wolbachia* genome sequencing depths

To further validate the presence of *Wolbachia* infections in *Anopheles* mosquitoes and provide more in-depth sequence analysis we undertook whole-genome sequencing.*Wolbachia* genome assemblies incorporated a minimum of 96.7 million paired-end (PE) reads per sample (from 160.9 million total) for *An. demeilloni* (*w*AnD) and 53.4 million PE reads (of 94.4 million total) for *An. moucheti* (*w*AnM). Additionally, we attempted to sequence *Wolbachia* genomes of one *An. coluzzii* from Ghana and five *An. gambiae* from the DRC that were *Wolbachia* positive by PCR^22^. Despite showing high sequencing depths against the mosquito host, there was little evidence of *Wolbachia* reads in any of these *An. gambiae* complex samples (**Fig. 5a, Supplementary table 4**). To further validate the presence of high-density *Wolbachia* strains within the two *Anopheles* species, we compared the genome coverage depth of the *w*AnD and *w*AnM genomes against 14 other *Wolbachia* strains sequenced with their hosts, using both sequencing data generated from this study as well as a selection of genomic sequencing data from different arthropods (see **Supplementary table 6** for accessions of the read data analysed and full results and **Supplementary table 7** for accessions of *Wolbachia* genomes). For *An. demeilloni* and *An. moucheti*, the average sequencing depth was comparable to arthropods with known *Wolbachia* strains (**Fig. 5b**, **Supplementary table 4**). In contrast, *An. coluzzii* and *An. gambiae* (including from Burkina Faso^15^) showed very low sequencing depth against *Wolbachia* genomes despite high sequencing depth against mosquito genomes which would be inconsistent with a maternally transmitted endosymbiont. These results clearly show that *An. demeilloni* and *An. moucheti* show comparable *Wolbachia* densities to mosquito species such as *Cx. quinquefasciatus* and *Ae. albopictus* which are known to contain resident *Wolbachia* strains in stable symbiotic associations. Simultaneously, our analysis further supports the concerns raised about the presence of a stable association between resident *Wolbachia* strains in *An. gambiae* and *An. coluzzii*.

**Fig. 5:**
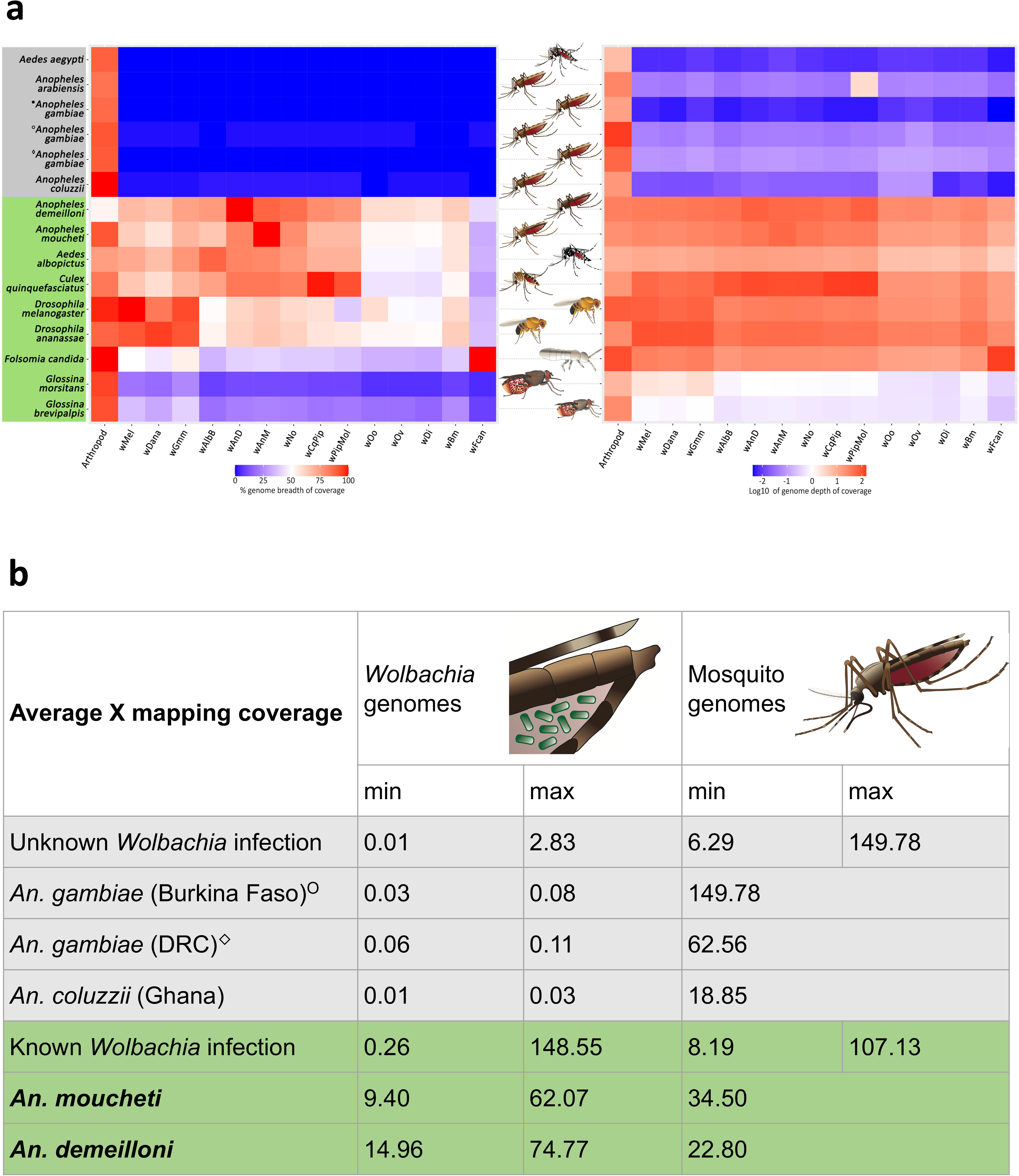
Breadth and depth of coverage of *Wolbachia* genomes. Insect hosts without a known *Wolbachia* strain are highlighted in grey, whilst those with a known *Wolbachia* infection are highlighted in green. Analysis includes *An. gambiae* from previously published studies (●), Burkina Faso - *Baldini et al. 2014* (○) and newly sequenced *An. gambiae* from the DRC (◊) and *An. coluzzii* from Ghana samples sequenced during our study. **a, Heatmap of coverage from published genome sequencing datasets after first mapping to the associated host genome and subsequently to a selection of *Wolbachia* genomes.** Shades of blue represent low values in either depth or breadth of coverage and shades of red represent high values. Samples from arthropods not known to contain *Wolbachia* have comparatively low depth and breadth of coverage for against *Wolbachia* genomes. A complete results table is available in Supplementary table 4. **b, Average mapping coverage of *Wolbachia* and host mosquito genomes.** The average minimum and maximum coverage are shown comparing *Anopheles* species and arthropods that have a known or unknown *Wolbachia* strain.

### *w*AnD and *w*AnM genome characteristics

These two newly sequenced genomes share key properties with other *Wolbachia* genomes, including genome size, predicted number of coding sequences and GC content (**Extended data Figs. 6-7, Supplementary table 5)**, and both our current and previously reported *Wolbachia* MLST and *wsp* gene analyses^23^ indicated both the *w*AnD and *w*AnM strains are supergroup B *Wolbachia* strains. To confirm this, we used a comparative Average Nucleotide Identity (ANI) analysis with 48 published *Wolbachia* genomes (**Supplementary tables 7-8**, **Fig. 6a)**. We also included an assembled *Wolbachia* genome that resulted from a recent large-scale computational study^29^ utilising *An. gambiae* genomes from the Ag1000G project^30^. The host species was subsequently classified as *An. species* A (here we have identified this species as *An. demeilloni*) and this genome shows close to 100% similarity to our assembled *w*AnD genome based on ANI analysis. The *w*AnD and *w*AnM strains cluster with other *Wolbachia* supergroup B strains (**Fig. 6b**) confirming the phylogenetic position indicated by MLST.

**Fig. 6:**
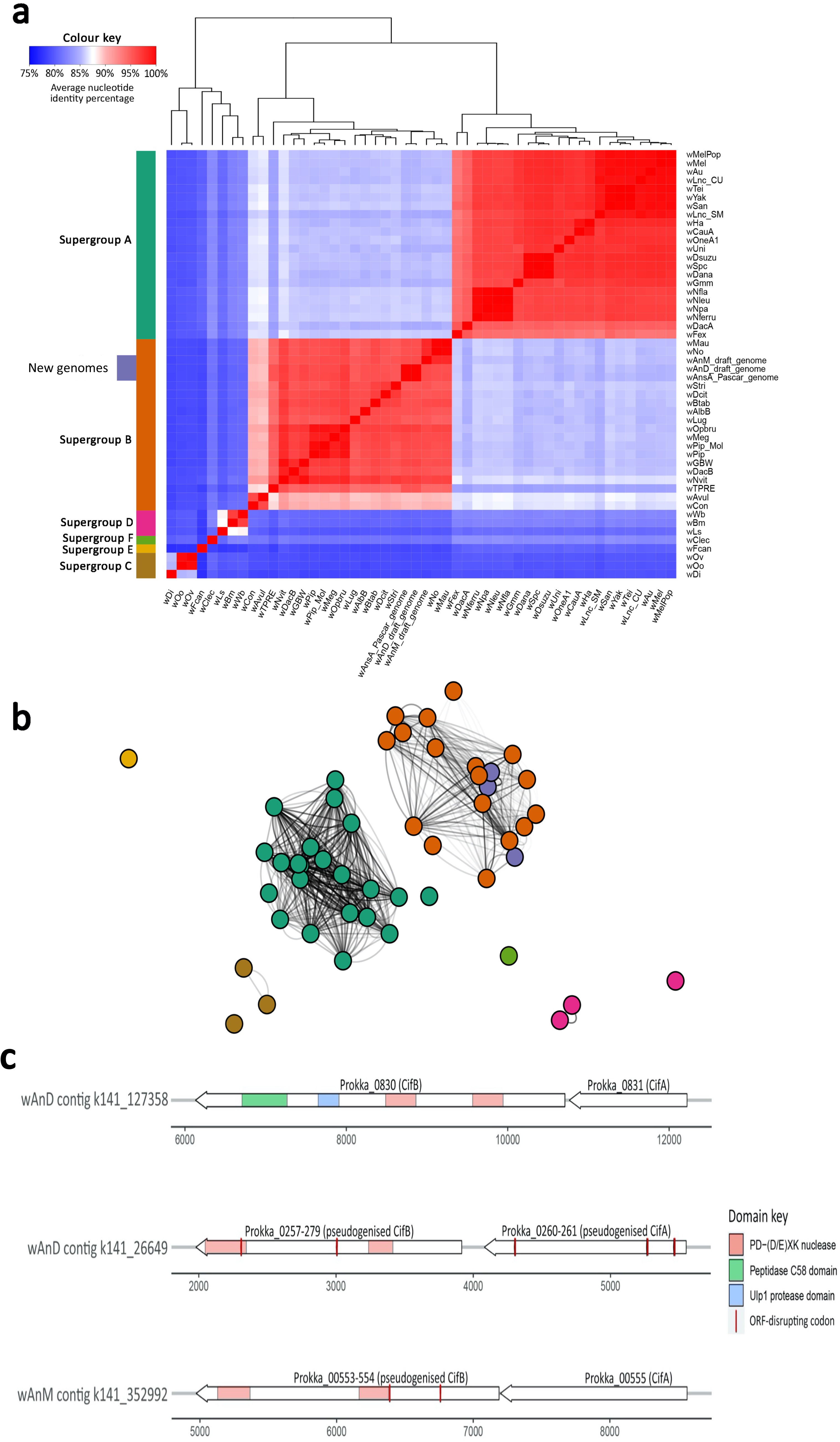
FastANI values, genome clustering analysis and *cif* genes. **a**, Heatmap showing the results of FastANI, comparing a total of 48 *Wolbachia* genomes against each other for similarity. High values represent close genetic similarity and a smaller phylogenetic distance, vice versa with low values, as shown by the colour key to the top left. The colour bar to the left of the heatmap indicates previously known clade organisation of the analysed *Wolbachia* species. **b,** Clustering analysis of the same *Wolbachia* genomes, with colour scheme preserved from the colour bar in panel **a**. The complete results matrix is available in supplementary table 8**. c,** Representation of the *cif* genes within the assembled *Wolbachia* genomes, with predicted protein domains overlaid. Each gene pair is drawn in relation to the contig they have been annotated on (x-axis, nucleotides). Domains were detected using the HHPred webserver

### CI genes are present in both *Wolbachia* strain genomes

. High prevalence rates, evidence of maternal transmission and high-density infections in wild mosquito populations indicated that both the *w*AnD and *w*AnM strains were likely CI-inducing strains containing CI factor (*cif*) genes associated with this phenotype in other *Wolbachia* strains^31–33^. *cifA* and *cifB* (and corresponding homologs) are neighbouring genes found across all CI-inducing strains and group into four monophyletic types^32,34^.

We identified two sets of *cif* gene homologs within the genome of *w*AnD, one of which however encodes multiple stop-codon and frame-shift interruptions (**Fig. 6c**). The predicted protein domains, as observed in previous studies^34^, included two PDDEXK nuclease domains, which are a consistent feature across all identified *cifB* genes. Compared to the previously assembled *Wolbachia* genome (from the host species classified previously as *An. species* A) from the Ag1000G project^29^, while it was noted that both pairs of *cif* genes were also present, the *cifB* gene of both pairs contained interruptions by either stop-codons or frame-shifts. In contrast to *w*AnD, the *w*AnM genome contained only one pair of *cif* genes, with the *cifB* gene interrupted with one stop codon and frame-shift **(Fig. 6c**).

## Discussion

Prior to this study, significant evidence showing that there is a stable association between *Anopheles* mosquitoes and endosymbiotic *Wolbachia* bacteria has been lacking^23^ despite an expanding number of studies which report amplification of *Wolbachia*-derived amplicons from *Anopheles* species. Criticism of previous studies is mainly based on their limitation to utilization of highly sensitive nested-PCR to amplify *Wolbachia* DNA from *Anopheles* isolates, which was extrapolated to indicate an endosymbiotic association^23,24^. To date, PCR-independent approaches that show the presence of live bacteria (like microscopy) rather than detection of DNA sequences have been lacking. The low infection frequencies and high variation in *Wolbachia* gene sequences of strains detected from *Anopheles* could be argued to be more consistent with environmental contamination rather than a stable bacterial endosymbiont that undergoes vertical transmission. Furthermore, variable gene sequences within the same mosquito species at a given location is inconsistent with well characterized *Wolbachia*-host endosymbiotic associations. Here we have addressed these concerns providing compelling evidence demonstrating that *An. moucheti* and *An*. *demeilloni* harbour high density maternally transmitted *Wolbachia* strains and show there is comparatively little evidence for stable native *Wolbachia* strains in the *An. gambiae* complex.

In our recent work, we reported the presence of potentially high-density infections in *An. moucheti* (n=1 from the DRC) and another *Anopheles* species which we have now resolved as *An. demeilloni*^22^. Here we expanded our screening of *Wolbachia* in these species including temporally and spatially spread sampling points. Our phylogeographic sequencing data show that the *w*AnM strain has an identical MLST and *wsp* gene profile in individuals from Cameroon to the original discovery in the DRC^22^. Furthermore, we present evidence of two *w*AnD strain variants (based on MLST profiles) present in both the DRC and Kenya. Taken together, these results show that both *Wolbachia* strains derived from the same host species span across large geographical areas which would be consistent with stably inherited CI-inducing strains. The prevalence rates in wild mosquito populations are also consistent with CI-inducing strains and is in direct contrast to the majority of studies that find a low prevalence rate of *Wolbachia* strains in *An. gambiae*, which is unusual for a maternally transmitted endosymbiont. Further studies are needed to determine whether genetic diversity within the *An. moucheti* complex could be influencing *Wolbachia* prevalence rates and how *Wolbachia* strain variation relates to genetic divergence within the *An. moucheti* complex, as indicated by our *COII* and corresponding *wsp* phylogenetic analysis. Interestingly, sequencing of the *w*AnM genome revealed an interrupted *cifB* gene which could also be indicative of variation in the levels of CI being induced by this strain.

To demonstrate the presence of live bacteria within the mosquito host, we also provide microscopic data showing intact *Wolbachia* cells in *Anopheles* ovaries using FISH. We show heavily infected ovarian follicles which are comparable to stable infection in the germline of naturally or artificially infected *Aedes* mosquitoes^35,36^. The punctate infection can be seen within the nurse cells that surround the oocyte which is often seen in *Wolbachia* infections in Diptera. These heavy ovarian infections are in contrast to the low levels of *Wolbachia* observed in *An. coluzzii* or our previous attempts to artificially infect *An. gambiae*, where small punctate infections were seen proximal to the follicular epithelium^17,37,38^. Our microscopy analysis found no evidence of infection in other mosquito tissues but this is likely explained by the lower prevalence rate and density of the *w*AnM strain in these tissues.

The density of *Wolbachia* strains in the *An. gambiae* complex and *An. funestus* s.s. are mostly reported at threshold levels of detection requiring nested PCR and providing only incomplete MLST profiles^16,18,19^. Furthermore, the inability to amplify and sequence the *wsp* gene also raises concerns given it is a commonly used marker for strain typing and is approximately 10 times more variable in its DNA sequence than the *16S rRNA* gene ^39^. In contrast, our qPCR and strain typing results presented here on larger cohorts of *An. moucheti* and *An. demeilloni* re-enforce that the *w*AnM and *w*AnD strains are present at significantly higher densities. In addition, the inability to find *Wolbachia* reads using microbiome sequencing in nested-PCR positive individuals raises concerns about the validity of this assay, which has been commonly used to report detection of *Wolbachia* infections in *Anopheles* mosquitoes^16–21^. A recent study using *16S rRNA* gene sequencing of nested PCR positive *An. coluzzii* from Burkina Faso found only one mosquito with 42 *Wolbachia* reads comprising 0.04% of relative abundance of the microbiome^40^. In comparison, our microbiome analysis shows that when present both the *w*AnM and *w*AnD strains dominate the microbiome which would be more consistent with a maternally transmitted endosymbiont.

Finally, evidence for high-density *Wolbachia* infections within these two *Anopheles* mosquito species is further confirmed by the assembly of near-complete genomes. In addition to this, read depths against the assembled genomes were comparable to those of other arthropods with known *Wolbachia* infections. We used a range of diverse *Wolbachia* strains as a scaffold and mapped reads from both published and our sequencing data sets. A high genome depth and coverage for both *w*AnM and *w*AnD *Wolbachia* genomes was seen even after sequencing through the more abundant host reads, with figures comparable to high-density *Wolbachia* infections seen in *Drosophila* species. This is in stark contrast to all *An. gambiae* complex sequencing data sets analysed, where the very low coverage is comparable to insects known not to harbour natural *Wolbachia* strains, and mapped reads likely to just represent background noise^15^.

Our reported high-density strains that localize in the germline appear desirable for vector control. The two genes responsible for *Wolbachia*-induced sperm modification and rescue (*cifA* and *cifB*) resulting in the CI phenotype were previously identified as part of prophage regions^32,41,42^ and our genome analysis provides strong evidence for the presence of *cif* gene homologs^43^. The induction of CI would be consistent with both high prevalence rates in wild mosquito populations and maternal transmission and would be a desirable feature for transinfection into other medically relevant *Anopheles* vector species. We microscopically observed a potentially higher *Wolbachia* density in ovaries compared to an *An. stephensi* transinfected line, suggesting the native strains are well adapted to their *Anopheles* hosts. While we did not observe *Wolbachia* in other tissues with microscopy, our qPCR data indicate somatic infection in some individuals. Whether the presence of these two high density *Wolbachia* strains would affect *Plasmodium* infection remains to be determined, but there are reports that lower density strains are correlated with *Plasmodium* inhibition^17,18^. In *Aedes* systems there is a positive relationship between *Wolbachia* density and viral interference but the role of density is less clear for *Wolbachia*-*Plasmodium* interactions^11,44^. Here we present robust data demonstrating for the first time high density *Wolbachia* strains naturally residing in *Anopheles* species which could potentially induce desirable phenotypes that make these strains stand out candidates for biocontrol strategies. Further characterization of the *w*AnM and *w*AnD strains in their ability to inhibit *Plasmodium* will provide the basis for their use in strategies to impact malaria transmission in wild mosquito populations.

## Methods

### Study sites, collection methods and historical sample collections

A variety of sampling methods were used to generate new mosquito collections in selected study sites in addition to analysis of historical DNA samples. *Anopheles* adult collections were undertaken in Olama Village (3.4125, 11.28416), Cameroon in June-July 2019 **(Supplementary table 9)** as this location has previously shown a high abundance of *An. moucheti*^45^. Human landing catches (HLCs) were undertaken between 19:00 and 06:00 for a total of 13 nights. In total, 104 Person/Trap/Nights were conducted, with 52 indoors and 52 outdoors. Trained volunteers were stationed at each house, with one individual inside and another outside. Participants exposed their legs and were provided with a flashlight. All mosquitoes that landed on exposed legs were collected in clear tubes and sealed with cotton wool. Tubes were organised into cotton bags labelled by hour, house number and location (indoors/outdoors). To reduce individual attraction bias, participants were rotated between houses for each night of collection, and halfway through each collection night the two volunteers at each house swapped places. All collection bags were transported from the field back to the Organisation de Coordination pour la lutte contre les Endémies en Afrique Centrale (Yaoundé, Cameroon) for morphological identification using keys^27^. Dead *An. moucheti* females were either stored in 100% absolute ethanol for subsequent PCR-based molecular analysis or in 100% acetone after removal of legs and wings to undergo FISH. Early generation colonisation was performed at OCEAC and later at LSHTM.

Larval sampling was undertaken in Lwiro (−2.244097, 28.815232), a village near Katana in the Democratic Republic of the Congo (DRC) in March 2019 to supplement existing mosquito DNA samples resulting from a 2015 collection containing a high abundance of *An. species* A individuals^46^. Larvae were collected and colonisation was performed at CRSN/LWIRO and later LSHTM. Morphological identification on adult females was independently carried out at LSHTM and CRSN/LWIRO (DRC) following keys^3,34^. Historical DNA samples of *An. species* A were also analysed from an area of Western Kenya^28^.

### DNA extraction and molecular mosquito species identification

Genomic DNA from whole bodies or dissected body parts (head-thorax and abdomens) were individually extracted using QIAGEN DNeasy Blood and Tissue Kits according to manufacturer’s instructions. DNA extracts were eluted in a final volume of 100 μL and stored at −20°C. To confirm species identification, a sub-set of individuals from all locations were subject to Sanger sequencing and phylogenetic analysis of ITS2^47^ and *COII*^48^ PCR products to enable greater differentiation of specimens. Sanger sequencing of PCR products was carried out as previously described ^22^ (sequences are listed in **Supplementary table 1**). To generate a rapid method for confirming mosquito species, ITS2 sequences for both *An. moucheti* and *An. demeilloni* were aligned **(Extended data Fig. 1)** and used to design species-specific qPCR assays (**Extended data Fig. 2)**. Forward and reverse primer sequences to amplify a fragment of the *An. moucheti* ITS2 were 5’-GTCGCAGGCTTGAACACA-3’ and 5’-ACTGTACCGCCTTACCATTTC-3’ respectively. Forward and reverse primer sequences to amplify a fragment of *An*. *demeilloni* ITS2 were 5’-GCTTAAGGCAGGTAAGGCGA-3’ and 5’-CGGTGTTAGAAGGCTCCGTT-3’ respectively. qPCR reactions were prepared using 5 μL of FastStart SYBR Green Master mix (Roche Diagnostics) with a final concentration of 1μM of each primer, 1 μL of PCR grade water and 2 μL template DNA, to a final reaction volume of 10 μL. Prepared reactions were run on a Roche LightCycler® 96 System for 15 minutes at 95°C, followed by 40 cycles of 95°C for 5 seconds, 60°C for 5 seconds and 72°C for 10 seconds. Amplification was followed by a dissociation curve (95°C for 10 seconds, 65°C for 60 seconds and 97°C for 1 second) to ensure the correct target sequence was being amplified.

### *Wolbachia* detection, quantification and confirmation of strain types

*Wolbachia* detection and quantification was undertaken targeting the conserved *Wolbachia 16S rRNA* gene^18^. BLAST analysis was first performed on previously generated *Wolbachia 16S rRNA* sequences for the *w*AnM and *w*AnD (previously known as *w*Ansa) strains of *Wolbachia*^22^ to confirm no sequence variability would influence primer binding. To estimate *Wolbachia* density across multiple *Anopheles* species, DNA extracts were added to Qubit™ DNA High Sensitivity Assays (Invitrogen) and total DNA was measured using a Qubit 4 Fluorometer (Invitrogen). A synthetic oligonucleotide standard (Integrated DNA Technologies) was used to calculate *16S rDNA* gene copies per μL using a ten-fold serial dilution^25^. *16S* rDNA gene real-time qPCR reactions were prepared using 5 μL of QIAGEN QuantiNova SYBR Green PCR Kit, a final concentration of 1μM of each primer, 1 μL of PCR grade water and 2 μL template DNA, to a final reaction volume of 10 μL. Prepared reactions were run on a Roche LightCycler® 96 System for 15 minutes at 95°C, followed by 40 cycles of 95°C for 15 seconds and 58°C for 30 seconds. Amplification was followed by a dissociation curve (95°C for 10 seconds, 65°C for 60 seconds and 97°C for 1 second) to ensure the correct target sequence was being amplified. Each mosquito DNA extract was run in triplicate alongside standard curves and NTCs and PCR results were analysed using the LightCycler® 96 software (Roche Diagnostics). Multilocus strain typing (MLST) was undertaken to characterize *Wolbachia* strains using the sequences of five conserved genes as molecular markers to genotype each strain^49^. PCR reactions and Sanger sequencing of PCR products were carried out as previously^22^. Sequencing analysis was carried out in MEGA X^50^ with consensus sequences used to perform nucleotide BLAST (NCBI) database queries, and for *Wolbachia* gene searches against the Wolbachia MLST database (http://pubmlst.org/wolbachia). Sanger sequencing traces from the *wsp* gene were also treated in the same way and analysed alongside the MLST gene locus scheme, as an additional marker for strain typing. All *Wolbachia* gene consensus sequences are listed in **Supplementary Table 1**).

### Phylogenetic analysis

Alignments were constructed in MEGA X^50^ by ClustalW to include relevant sequences highlighted through searches on the BLAST and *Wolbachia* MLST databases. Maximum Likelihood phylogenetic trees were constructed from Sanger sequences as follows. The evolutionary history was inferred by using the Maximum Likelihood method based on the Tamura-Nei model^51^. The tree with the highest log likelihood in each case is shown. The percentage of trees in which the associated taxa clustered together is shown next to the branches. Initial tree(s) for the heuristic search were obtained automatically by applying Neighbor-Join and BioNJ algorithms to a matrix of pairwise distances estimated using the Maximum Composite Likelihood (MCL) approach, and then selecting the topology with superior log likelihood value. The trees are drawn to scale, with branch lengths measured in the number of substitutions per site. Codon positions included were 1st+2nd+3rd+Noncoding. All positions containing gaps and missing data were eliminated. The phylogeny test was by Bootstrap method with 1000 replications. Evolutionary analyses were conducted in MEGA X^50^.

### Microbiome analysis

The microbiomes of selected individual mosquitoes were analysed using barcoded high-throughput amplicon sequencing of the bacterial *16S rRNA* gene (with library preparation and Illumina sequencing carried out commercially by Source BioScience, Cambridge, UK). Detailed methodology is provided in supplementary methods file.

### Fluorescence *in situ* hybridization (FISH)

Freshly dead adult female mosquitoes were fully submerged in 100% acetone after removal of all legs and wings. Whole mosquitoes were embedded in paraffin wax and sectioned at Liverpool Bio-Innovation Hub (University of Liverpool). The FISH protocol was conducted as previously reported^52^. Briefly, sections were deparaffinated with three 5-minute washes in 100% Xylene, one 5-minute wash in 100% EtOH and one 5-minute wash in 95% EtOH. Slides were then placed in 6% H_2_O_2_ and 80% EtOH for at least 4 days. Slides were washed with diH_2_O and 50ng of Wol3_Red (/5ATTO590N/TCCTCTATCCTCTTTCAATC) and 50ng of Wol4_Red (GAGTTAGCCAGGACTTCTTC/3ATTO590N/) were added to 500 ul of hybridisation buffer pre-heated to 37°C^53^. Buffer containing the probes was placed on the slide and slides were placed in a hybridisation chamber overnight at 37°C. Slides were washed once in 1x SSC (10mM DTT) for 15 mins, twice in 1x SSC (10mM DTT) for 15 mins at 55°C, twice in 0.5x SSC (10mM DTT) for 15 mins at 55°C, and finally, once in 0.5x SSC (10mM DTT) for 15 mins. Slides were again washed with diH_2_O and 2ul of DAPI in 200ul of 1xPBS was placed on the tissue for 8 minutes. Slides were washed with 1x PBS and slides were mounted with a drop of anti-fade. No probe and competition controls were undertaken. We also had a positive control which was *Cx. quinquefasciatus and Ae. albopictus* mosquitoes that harbour *Wolbachia*. Images were captured with a Revolve FL microscope (Echolab).

### Genome sequencing

Raw pair-ended reads from *An. gambiae* (n=4), *An. demeilloni* (n=3), *An. coluzzii* (n=1) and *An. moucheti* (n=1) were trimmed for Illumina Nextera adapter sequences using Trimmomatic^54^. Reads were also quality-trimmed with Trimmomatic to a minimum PHRED quality of 20 within a sliding window of 4, discarding reads that fell below a minimum length of 100 base-pairs. Subsequently, host mosquito reads were removed from the samples. As no reference genome exists for either *An. moucheti* or *An. demeilloni*, genome assemblies of *An. gambiae* (accession AgamP4), *An. funestus* (accession AfunF3), and *An. arabiensis* (accession AaraD1) were downloaded from VectorBase^55^. The trimmed pair-ended reads were mapped against the genome of *An. gambiae* using the BWA aligner with default settings (version 0.7.12-r1039)^56^. Unmapped reads were extracted from the alignment, and remapped against the genome of *An. funestus*, before remaining unmapped reads were extracted and remapped to the genome of *An. arabiensis*. Only reads that remained after this sequential remapping to three different *Anopheles* mosquito genomes were taken forward for *de-novo* genome assembly.

### Genome assembly

*De-novo* genome assembly was conducted using the program MEGAHit^57^ with default parameters, which utilises succinct de-brujin graphs for resource-efficient assembly of contigs from metagenomic data. This generated two sets of contigs from the two different mosquito species that were then analysed with MetaQUAST^58^ to identify microbial species present within the dataset. The closest *Wolbachia* genome of *D. simulans* strain Noumea (*w*No)^59^ was selected (NCBI accession no. CP003883.1). The *w*No genome was used to create a BlastN database and all contigs generated by MEGAHit^57^ were searched against the *w*No genome to identify contigs that are of likely *Wolbachia* origin within the two *Anopheles* species. These identified contigs were scaffolded against the ‘reference’ *Wolbachia* genome using the Mauve contig mover^60,61^.

Reads from the two mosquito datasets were remapped to their corresponding draft genome assembly with the BWA aligner^56^ using default settings and average read depth calculated for each contig using the program samtools depth^62^. Contigs that showed greater than one standard deviation from the average read depth were removed from the assembly. Subsequent to the removal of these contigs, the reads were remapped to the draft genome and subsequently used to improve the assembly using the program Pilon^63^. Pilon automatically detects for the presence of single nucleotide variants, or insertions/deletion events introduced during the assembly process. This was repeated a total of three times, where no further insertion/deletions were detected.

### Genome annotation and comparisons to existing genomes and sequence data

Annotation of both *Wolbachia* genomes was performed using the program PROKKA^64^ using default settings. This annotation was used to check for genome completeness using CheckM^65^ and identification of cytoplasmic incompatibility factor (*cif*) genes. The assembled genome sequence was further used for comparison against existing genomes using the program FastANI^67^, and further used for comparing read coverage and depth against a selection of other *Wolbachia* species. Detailed methodology is provided in supplementary methods.

## Statistical analysis

Normalised qPCR *Wolbachia 16S* rRNA gene copies per μL were compared using unpaired t-tests in GraphPad Prism 7.

## Ethical approval

Ethical approval for undertaking HLCs in Cameroon was obtained from the LSHTM ethics committee (reference no. 16684) in addition to local ethical approval (clearance no. 2016/01/685/CE/CNERSH/SP) delivered by the Cameroon National Ethics (CNE) Committee for Research on Human Health). Informed consent was gained from all volunteers prior to commencement of sampling and all volunteers were provided with malarial chemoprophylaxis.

## Data availability

All data supporting the findings of this study are available within the article, as Supplementary Information and raw qPCR data is available at https://doi.org/10.17605/OSF.IO/AHNB6. Raw sequencing data has been uploaded to NCBI under BioProject PRJNA642000, accession numbers SRR12095496 through to SRR12095498, and SRR12729562. Sanger sequencing data is available with accession numbers XXXX – XXXX.

## Acknowledgements

The authors would like to thank Ralph Harbach of the Natural History Museum (London, UK) for independent expert morphological identification of mosquito samples. TW and CLJ were supported by a Sir Henry Dale Wellcome Trust/Royal Society fellowship awarded to TW (101285): http://www.wellcome.ac.uk;https://royalsociety.org. GLH was supported by the BBSRC (BB/T001240/1), the Royal Society Wolfson Fellowship (RSWF\R1\180013), NIH grants (R21AI124452 and R21AI129507), UKRI (20197), and the NIHR (NIHR2000907). GLH is affiliated to the National Institute for Health Research Health Protection Research Unit (NIHR HPRU) in Emerging and Zoonotic Infections at University of Liverpool in partnership with Public Health England (PHE), in collaboration with Liverpool School of Tropical Medicine and the University of Oxford. GLH is based at LSTM. The views expressed are those of the author(s) and not necessarily those of the NHS, the NIHR, the Department of Health or Public Health England. GLH and EH are also jointly funded by the BBSRC (V011278/1). SH was supported by the LSTM Director’s Catalyst Fund award.

## Author contributions.

T.W. acquired funding, supervised mosquito field collections, undertook laboratory analysis of samples, provided overall supervision alongside G.H. and co-wrote the first draft. S.Q. undertook genome assemblies and genome analysis and co-wrote the first draft. C.L.J. undertook laboratory analysis of samples, sanger sequence analysis, microbiome analysis, supervision of field collections and co-wrote the first draft. J.B. undertook mosquito colonisation and provided samples from the DRC. V.D. contributed to mosquito collection, colonisation and laboratory analysis of samples. R.B. contributed to mosquito collection and colonisation. M.K. contributed to laboratory analysis of samples and microbiome analysis. L.A.M. contributed to laboratory analysis of samples and microbiome analysis. A.G. contributed to mosquito collection and laboratory analysis of samples. E.A.H. contributed to microbiome analysis. E.R.A. undertook FISH. C.C-U. contributed to figure preparation. S.H. contributed to FISH. C.B. provided reagents and support for mosquito colonisation in the DRC. J.C.S. and N.F.L. contributed mosquito DNA samples from Kenya. S.C.W. contributed to *Wolbachia* genomic sequencing analysis. C.A.N. provided logistical support and supervision of fieldwork in Cameroon. E.H. supervised *Wolbachia* genome sequence analysis and co-wrote the first draft. G.L.H. acquired funding, supervised FISH and provided overall supervision alongside T.W. and co-wrote the first draft.

## Materials & Correspondence.

Thomas Walker and Grant Hughes

## Supplementary information

### Extended Data Figure legends

**Extended data Fig. 1 Alignment of ITS2 sequences and location of species-specific primers.**

**Extended data Fig. 2 ITS2 species specific qPCR fluorescence targeting *An. demeilloni* (a) and *An. moucheti moucheti* (b).** Inset = dissociation curves to ensure the correct target sequence was being amplified. dem = *An. demeilloni, mou* = *An. moucheti moucheti, gam = An. gambiae* s.s.*, col = An. coluzzii*.

**Extended data Fig. 3. Images of ‘ *An. species* A’.** a=adult female, b=wing of adult female, c=adult male. d=larvae. Independent morphological identification by three individuals using keys confirmed this species is *An. demeilloni*.

**Extended data Fig. 4. Controls for FISH.** *Wolbachia*-infected *Cx. quinquefasciatus* samples used as a positive control A-D. *An. moucheti* competitive control E-H*. An. moucheti* no probe control I-L (Scale bars 90μM in A-L).

**Extended data Fig. 5 Microbiome individual sample relative taxonomic abundance barplots.** The relative taxonomic abundance barplots for each sample, as visualised using the qiime taxa barplot command within QIIME2^66^. Sample groups with metadata are as detailed in the table and the legend details the level 7 classification of the 20 most abundant ASVs across all samples. Samples are arranged by group, then by descending % *Wolbachia*. The overwhelming dominance of *Wolbachia* within the microbiome of *An. demeilloni* and *An. moucheti* samples in the *Wolbachia* positive groups can be seen. In addition, the presence of high numbers of *Wolbachia* reads across different years for *An. demeilloni* (groups B and C), and in both the abdomen and head-thorax in *An. moucheti* (groups C and D) is shown. The maternal transmission of *Wolbachia* is demonstrated in the F1 *An. moucheti Wolbachia* positive samples within group H. The diversity of microbes present in these mosquitoes when *Wolbachia* is absent can also been seen (groups A, F, G and H).

**Extended data Fig. 6 Similarity plot of the *w*AnD genome compared against a selection of other *Wolbachia* genomes.** The BLAST Ring Image Generator (BRIG) program was used to analyse the percentage identity of the *w*AnD genome against 5 other *Wolbachia* genomes, including the *w*AnD genome itself. Each coloured ring from the centre represents a different *Wolbachia* genome as represented on the key to the top right of the image, with the saturation of colour at certain coordinates of the circle representing how conserved that region of the *w*AnD genome is when compared against the target *Wolbachia* genome.

**Extended data Fig. 7 Similarity plot of the *w*AnM genome compared against a selection of other *Wolbachia* genomes.** See information for Extended data Figure 6.

**Extended data Fig. 8 Heatmap representing depth of coverage for the assembled *w*AnM and *w* AnD genomes within 10kbp windows.** Contigs for both genomes were first concatenated into one long assembly, before being separated into 10kbp-long windows. Sequencing data from individual samples were then mapped against the genome, and sequencing depth for each 10kbp window then calculated. Each row represents a single sample that has been aligned to one of the two genomes *w*AnM or *w*AnD, with intensity of red indicating the depth of sequencing as shown by the key to the right.

### Supplementary tables

**Supplementary table 1. Additional Sanger sequencing sample details for *w*AnM-infected *An. moucheti* and *w*AnD-infected *An. demeilloni* with their associated GenBank accession numbers.** The location and year of collection, sample codes and the sequenced gene fragment is shown in addition to the GenBank accession number.

**Supplementary table 2. *Wolbachia* density of the *w*AnM and *wAnD* strains.** Mean *Wolbachia 16S rRNA* gene copies/ng DNA for *Wolbachia*-infected mosquito DNA extracts. *approximately 200 eggs were pooled prior to extraction. CAM = Cameroon, DRC = Democratic Republic of Congo, KEN = Kenya.

**Supplementary table 3. *w*AnM and *w*AnD *Wolbachia* strain *WSP* typing.** The *wsp* sequence for *w*AnM-VAR2 had 1 nucleotide difference to allele number 322 (CM = closest match).

**Supplementary table 4. Read depth analysis results.** Full results from read mapping analysis of other DNA-sequencing datasets against a selection of 14 *Wolbachia* genomes. For each of the analysed DNA-sequencing datasets, represented by the corresponding SRA number, mapping metrics for both depth and breadth of coverage against the originating arthropod genome, as well as the selection of 14 different *Wolbachia* genomes, was included.

**Supplementary Table 5. General characteristics of the *w*AnD and *w*AnM genomes.**

**Supplementary table 6. CheckM results.** Full results from CheckM with regards to genome completeness of the two assembled genomes from this study, as well as a selection of other *Wolbachia* genomes.

**Supplementary table 7. *Wolbachia* genomes used for comparison.** Existing *Wolbachia* genomes used in this study for comparison against the assembled genomes.

**Supplementary table 8. FastANI results.** Full results matrix from FastANI with regards to genome similarity between multiple *Wolbachia* genomes. Numbers are on a percentage similarity scale, as determined by FastANI, with 100% indicating exact similarity, and vice versa.

**Supplementary table 9. *Anopheles* species collected from Olama Village, Cameroon using human landing catches in 2019.**

