## Supplementary figures and images for "Genomic and microscopic evidence of stable high density and maternally inherited *Wolbachia* infections in *Anopheles* mosquitoes"

### Extended data Figure 1

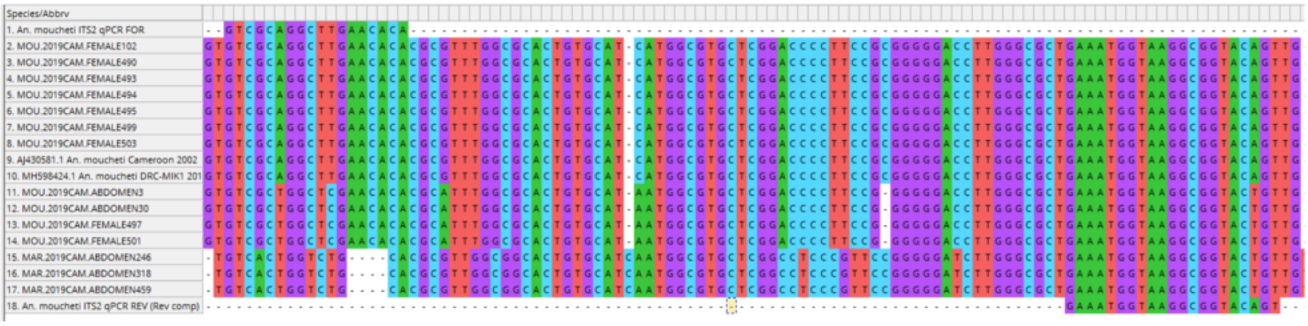

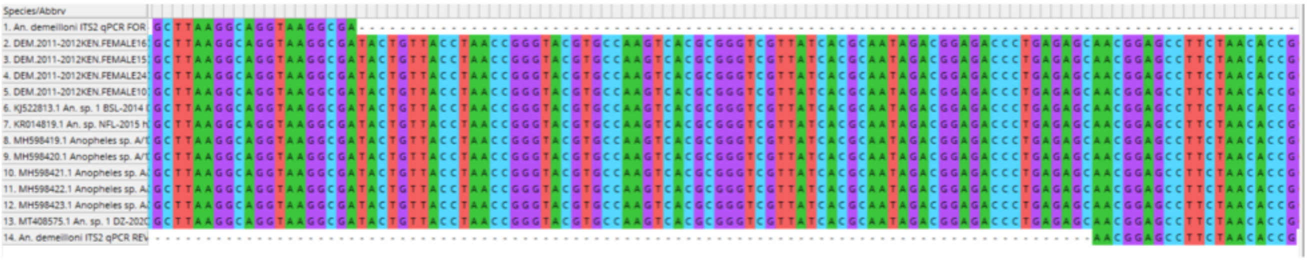

### Extended data Figure 2

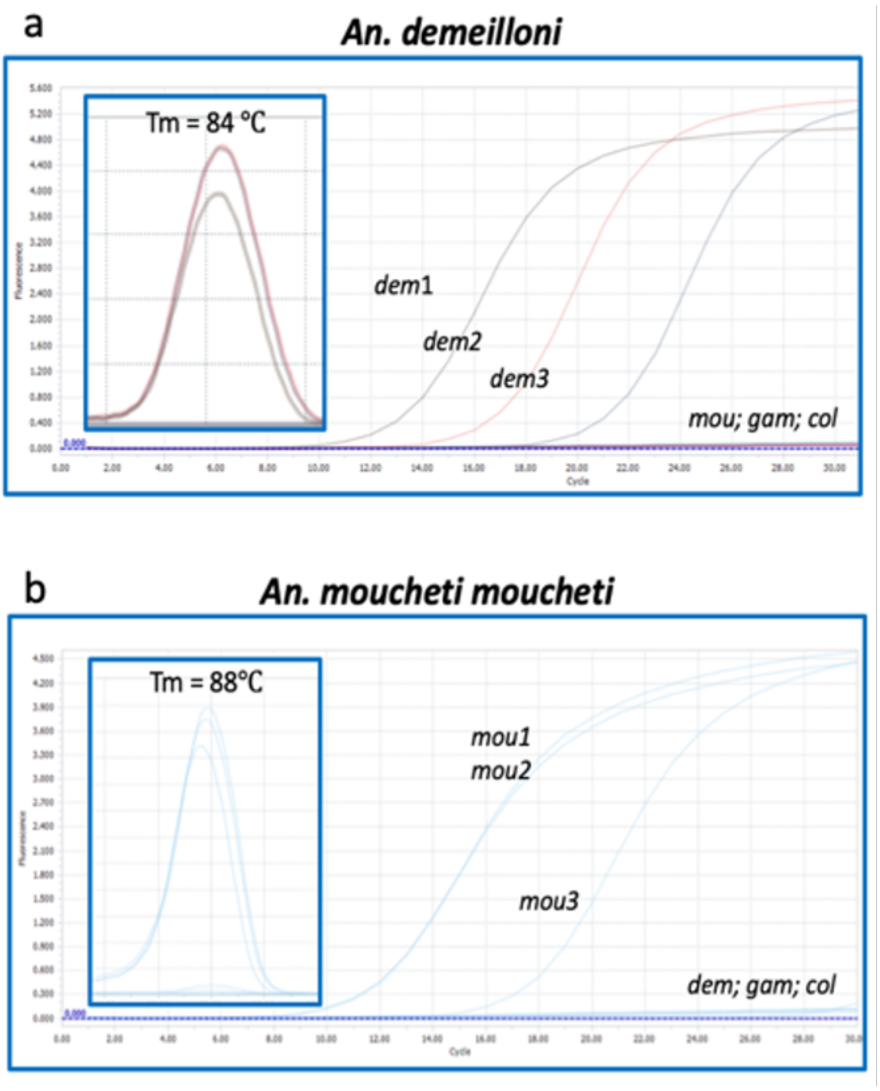

### Extended data Figure 3

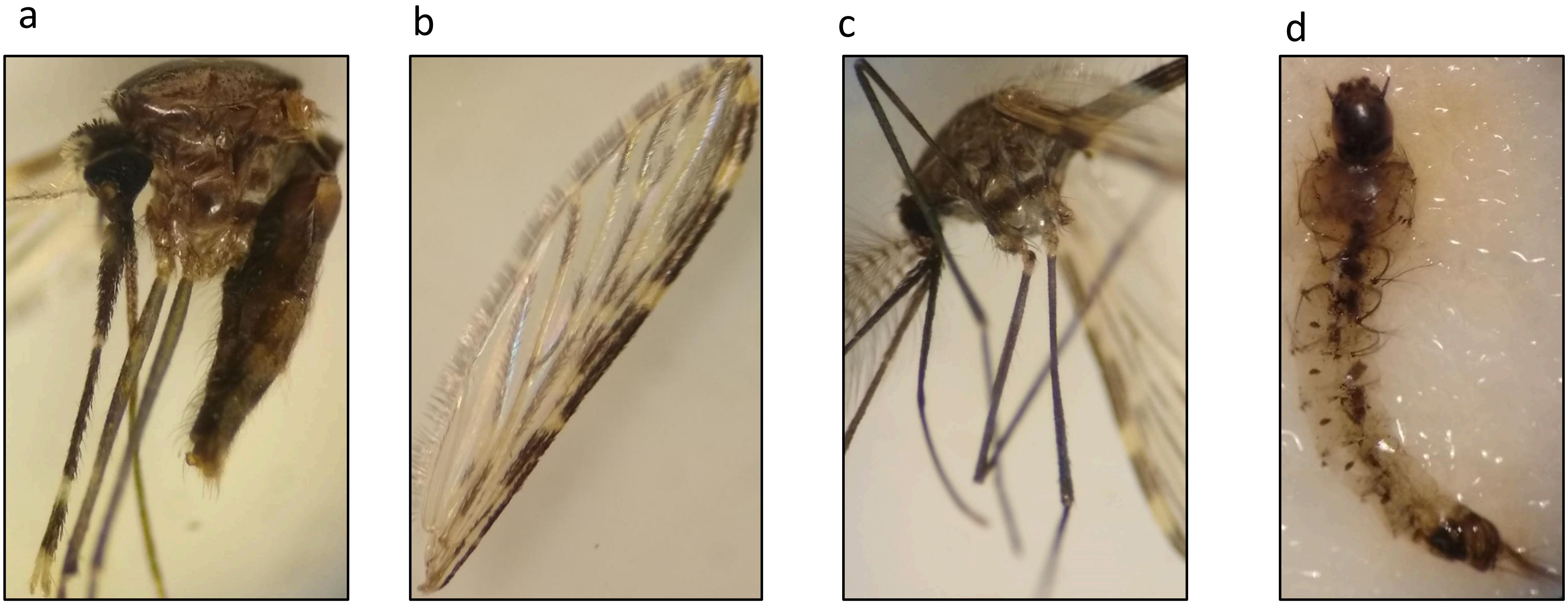

### Extended data Figure 4

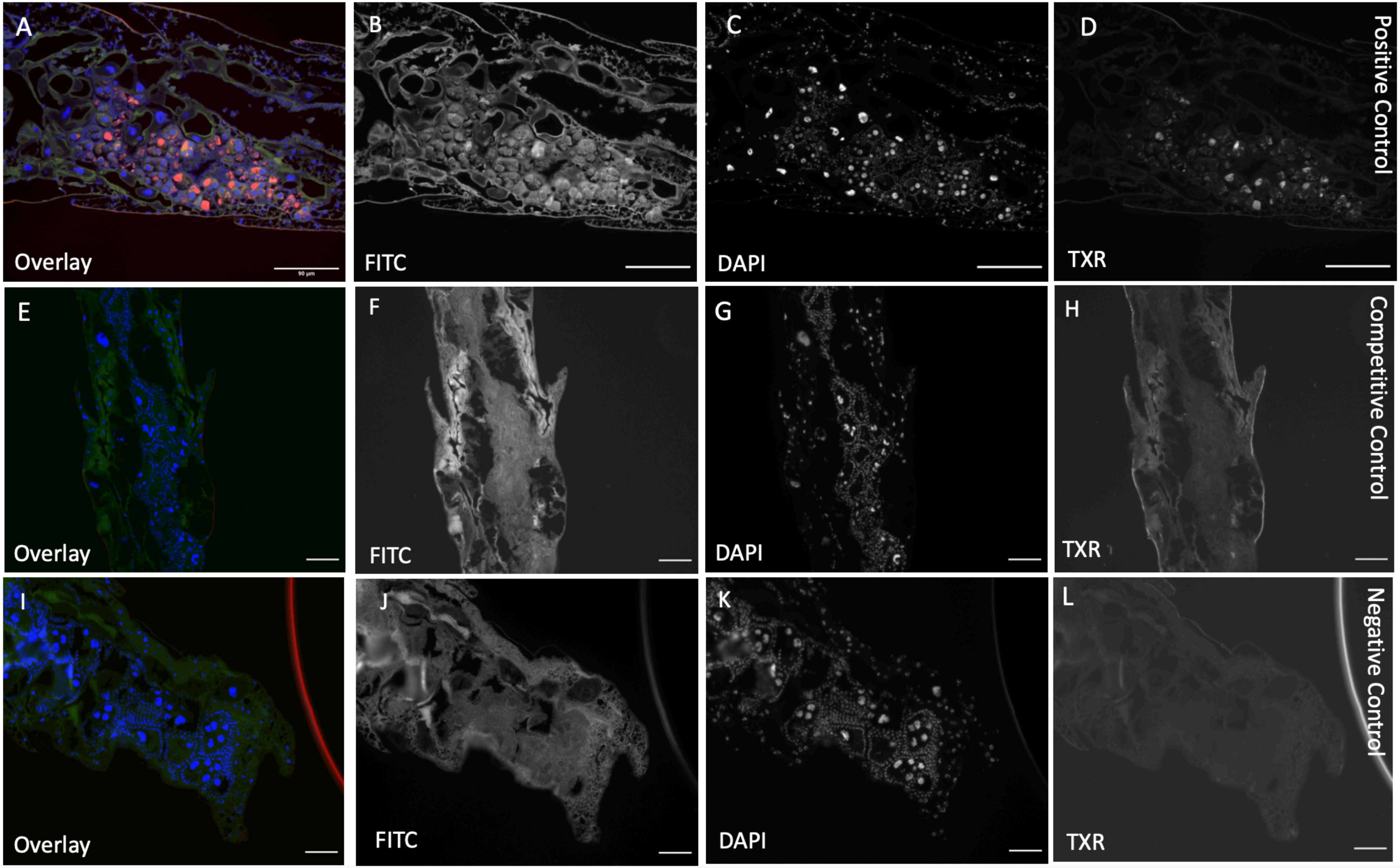

### Extended data Figure 6

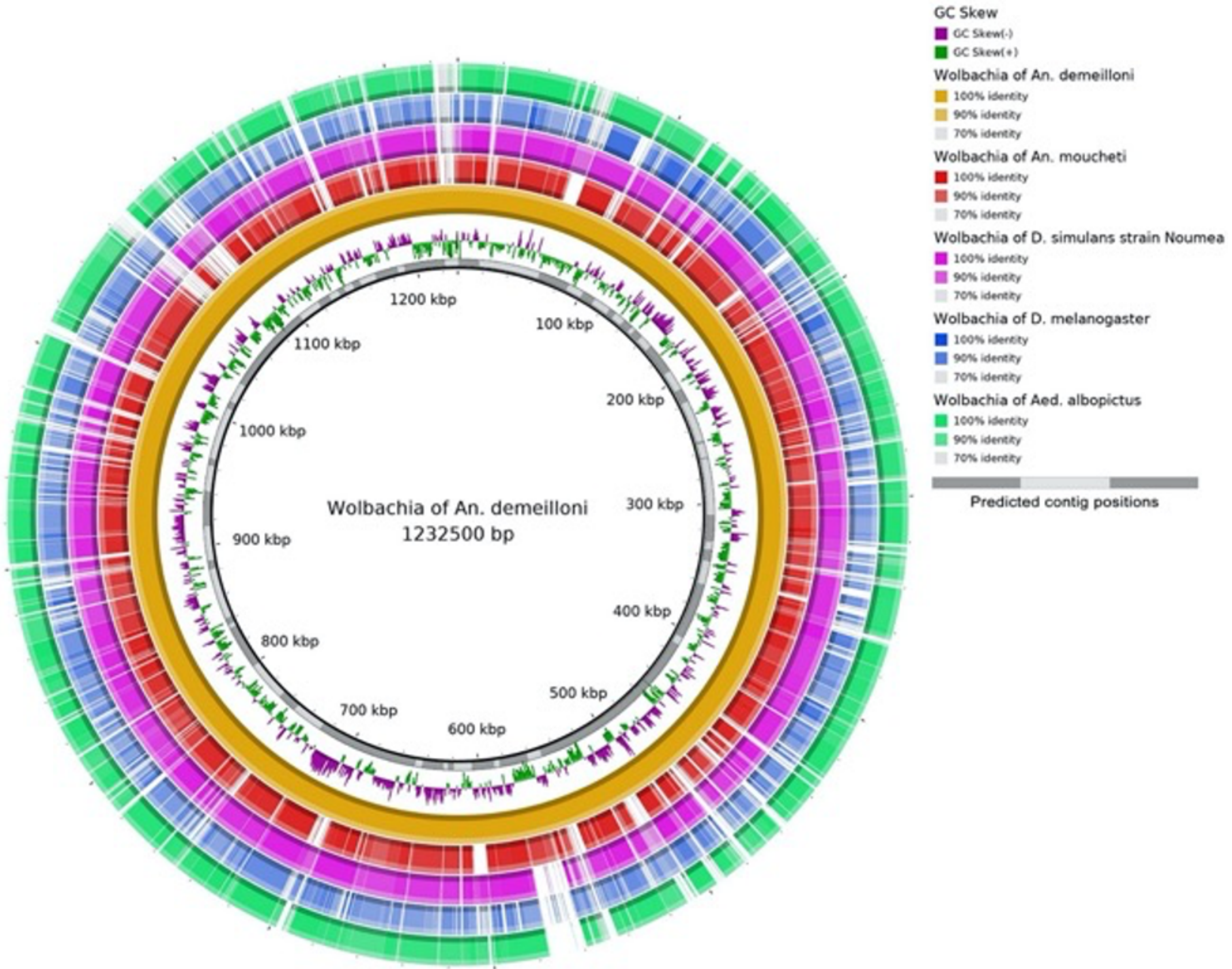

### Extended data Figure 7

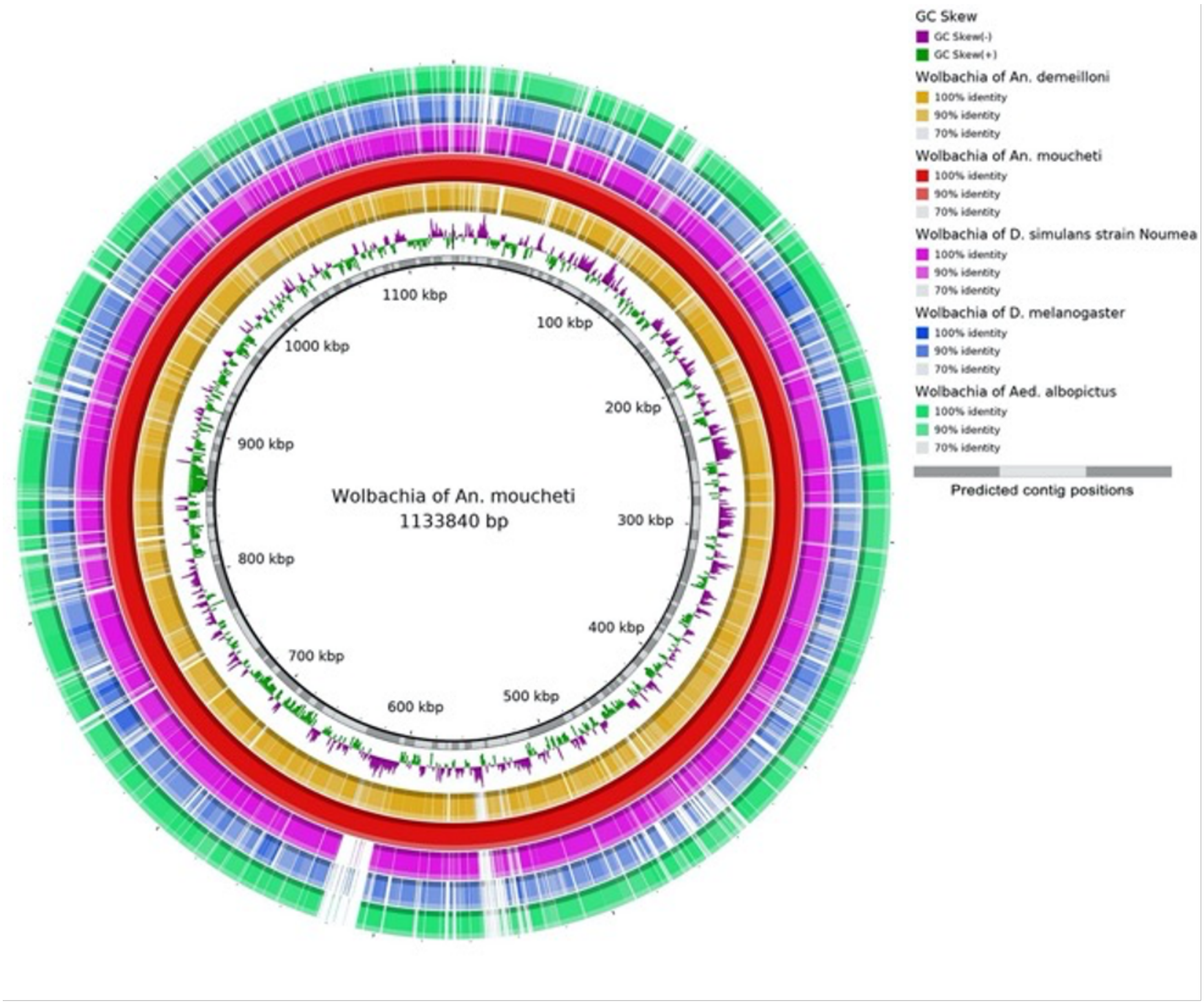

### Extended data Figure 8

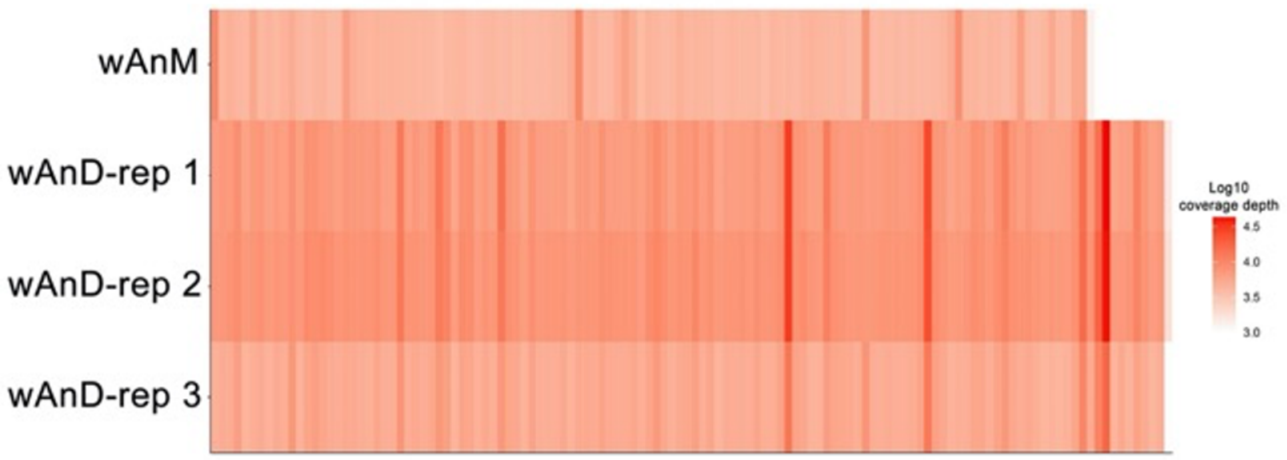
