## Extended data Figure 5 for "Genomic and microscopic evidence of stable high density and maternally inherited *Wolbachia* infections in *Anopheles* mosquitoes"

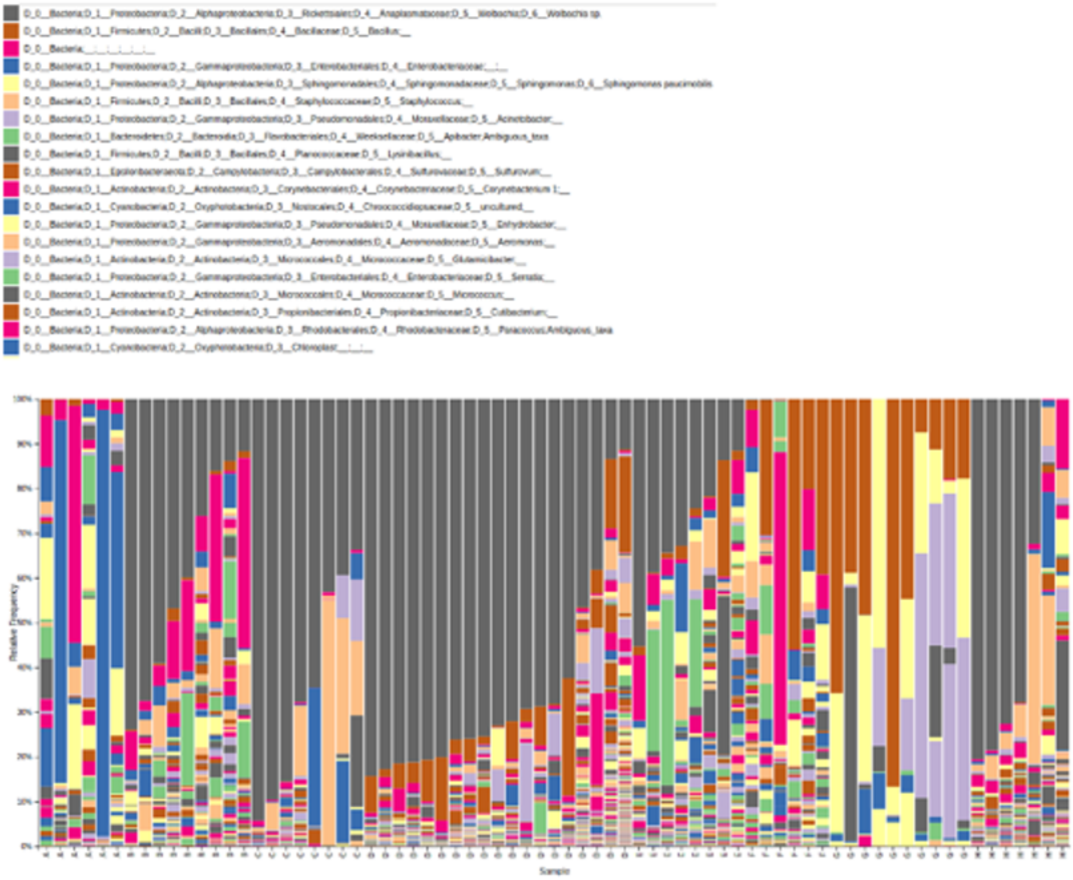


| Group-id | Species | Country | Body-part | Year | Number | Wolbachia-status |
|---|---|---|---|---|---|---|
| A | *An-demeilloni* | DRC | whole | 2015 | 6 | Negative |
| B | *An-demeilloni* | DRC | whole | 2015 | 9 | Positive |
| C | *An-demeilloni* | DRC | whole | 2019 | 8 | Positive |
| D | *An-moucheti* | Cameroon | abdomen | 2019 | 19 | Positive |
| E | *An-moucheti* | Cameroon | head-thorax | 2019 | 8 | Positive |
| F | *An-moucheti* | Cameroon | head-thorax | 2019 | 6 | Negative |
| G | *An-moucheti* | Cameroon | whole | 2019 | 10 | Negative |
| H | *An-moucheti (F1)* | Cameroon | whole | 2019 | 7 | Positive (5) and Negative (2) |
